# Genome Report: *De novo* assembly of a high-quality reference genome for the Horned Lark (*Eremophila alpestris*)

**DOI:** 10.1101/811745

**Authors:** Nicholas A. Mason, Paulo Pulgarin, Carlos Daniel Cadena, Irby J. Lovette

## Abstract

The Horned Lark (*Eremophila alpestris*) is a species of small songbird that exhibits remarkable geographic variation in appearance and habitat across an expansive distribution. While *E. alpestris* and related species have been the focus of many ecological and evolutionary studies, we still lack a highly contiguous genome assembly for horned larks and related taxa (Alaudidae). Here, we present CLO_EAlp_1.0, a highly contiguous assembly for horned larks generated from blood samples of a wild, male bird captured in the Altiplano Cundiboyacense of Colombia. By combining short-insert and mate-pair libraries with the ALLPATHS-LG genome assembly pipeline, we generated a 1.04 Gb assembly comprised of 2708 contigs with an N50 of 10.58 Mb and a L50 of 29. After polishing the genome, we were able to identify 94.5% of single-copy gene orthologs from an Aves data set and 97.7% of single-copy gene orthologs from a vertebrata data set, indicating that our *de novo* assembly is near complete. We anticipate that this genomic resource will be useful to the broader ornithological community and those interested in studying the evolutionary history and ecological interactions of a widespread, yet understudied lineage of songbirds.

## Introduction

The Horned Lark (*Eremophila alpestris*) is a widespread species of songbird that occupies grasslands, tundras, deserts, and other sparsely vegetated habitats on five continents (Beason 1995). As is characteristic of most species in the family Alaudidae, *E. alpestris* is a terrestrial species that nests on the ground and relies on camouflage to avoid predation by avian predators (Donald *et al.* 2017). The Horned Lark has been studied extensively in terms of geographic variation and systematics (Behle 1942; Johnson 1972), population genetics (Drovetski *et al.* 2006, 2014; Mason *et al.* 2014; Ghorbani *et al.* 2019), physiological adaptations (Trost 1972), breeding biology (de Zwaan *et al.* 2019), and responses to human activity, such as agriculture (Mason and Unitt 2018) and wind energy (Erickson *et al.* 2014), among other focal areas. Despite extensive past and ongoing research involving *E. alpestris* and other alaudids, we lack a highly contiguous reference genome for the species and the family as a whole (but see (Dierickx *et al.* 2019)). Generating genomic resources for horned larks and related taxa will enable studies linking phenotypic and genetic variation (Kratochwil and Meyer 2015; Hoban *et al.* 2016), chromosomal rearrangements (Wellenreuther and Bernatchez 2018), and many other avenues of future genomic research for non-model organisms (Ellegren 2014).

Here, we describe CLO_EAlp_1.0, a new genomic assembly that we built with DNA extracted from a wild, male lark captured and from a demographically small and geographically isolated population near Toca, Boyacá, Colombia. We sampled this individual and population because it had high *a priori* likelihood of high homozygosity compared to larks elsewhere with much larger effective population sizes and variable patterns of connectivity to adjacent populations. To generate this de novo assembly, we used the ALLPATHS-LG pipeline (Butler *et al.* 2008; Gnerre *et al.* 2011). Given the lack of genomic resources currently available for Alaudidae, we hope this de novo assembly will inspire and facilitate future studies on the genomic biology of larks—a widespread, diverse lineage of songbirds.

## Methods

### Sample collection, DNA extraction, and sequencing

We captured a male *E. alpestris* (EALPPER07; NCBI BioSample SAMN12913182) approximately 170 km NE of Bogotá, Colombia near the town of Tocá on the shores of the Embalse de La Copa in the Altiplano Cundiboyacense of the Boyacá department (5.623299° N, 73.184156° W). This population is small and represents a subspecies (*E. a. peregrina*) that is geographically isolated from other populations of larks, the nearest population of which is in Oaxaca, Mexico. The Colombian subspecies of Horned Lark underwent a population bottleneck upon colonizing the high-elevation plateaus of the region and therefore has high homozygosity, which is preferable for *de novo* genome assembly. We collected blood from the brachial vein, from which we subsequently extracted genomic DNA with a Gentra Puregene Blood Kit (Qiagen, Hilden, Germany) following the manufacturer’s protocol. We confirmed the sex of the individual using PCR amplification (Chu *et al*. 2015). After running the sample on a 1% agarose gel to confirm the presence of high molecular weight DNA, we sent the extraction to the Cornell Weil Medical School, where they generated a 180 bp fragment library, a 3 kb mate-pair library and a 8 kb mate-pair library. We sequenced the 180 bp library across two lanes and combined the 3 kb and 8 kb mate-pair libraries on another lane of Illumina HiSeq 2500 to perform 100 bp paired-end sequencing.

### Genome assembly, polishing, and assessment

We assembled the genome with ALLPATHS-LG v52415 (Butler *et al.* 2008; Gnerre *et al.* 2011). We did not perform additional adapter removal or quality filtering with the short-insert 180 bp libraries because ALLPATHS-LG has built-in steps that remove low quality and adapter-contaminated reads (Butler *et al.* 2008). Once the initial assembly had finished, we aligned the short-insert and mate-pair libraries back to the assembly genome using bwa 0.7.17-r1188 (Li and Durbin 2009) and samtools v1.9 (Li *et al.* 2009) and then performed three iterations of scaffold polishing using pilon v1.22 (Walker *et al.* 2014) with default parameters. Once scaffold polishing had finished, we ordered and correspondingly renamed the scaffolds with respect to decreasing scaffold sizze using SeqKit v0.7.2 (Shen *et al.* 2016). We assessed the contiguity the *de novo* genome using QUAST v5.0.2 (Mikheenko *et al.* 2018) and estimated genome completeness with BUSCO v3 (Simão *et al.* 2015; Waterhouse *et al.* 2018) alongside HMMER v3.1b2 (Finn *et al.* 2011) and BLAST+ v2.7.1 (Camacho *et al.* 2009) to identify single-copy orthologous gene sets among birds and vertebrates.

### Mitochondrial Genome Assembly

We also assembled the mitochondrial genome for the same individual (EALPPER07) with NOVOplasty v3.7 (Hahn *et al.* 2013) using a ND2 sequence (GenBank Accession KF743558) from a previous study (Mason *et al.* 2014) as the initial seed to begin the assembly process.

### Data availability

Raw output from sequencing runs and the final assembly, CLO_EAlp_1.0, are available from NCBI (BioProject PRJNA575884). Short-fragment and mate-pair libraries are also available from the NCBI SRA (SUB6392689). Outputs from BUSCO and QUAST analyses are available from FigShare (doi:10.6084/m9.figshare.9956063;doi:10.6084/m9.figshare.9956042).

## Results and Discussion

Taken together, the three lanes of Illumina HiSeq 2500 sequencing generated 1.59 × 10^9^ total reads (~134x estimated coverage of a 1.2 Gb genome), including 5.45 × 10^8^ paired-end reads for the 180 bp short-insert libraries, 1.24 × 10^8^ paired-end reads for the 3 kb mate-pair library, and 1.27 × 10^8^ paired-end reads for the 8 kb mate-pair library. Following scaffold polishing, the finalized CLO_EAlp_1.0 assembly consisted of 2708 contigs that totaled 1.04 Gb. The largest contig was 31.81 Mb while the N50 was 10.58 Mb and L50 was 29 (Table 1). The average GC content was 42.23%, which is similar to other birds (Jarvis *et al.* 2014; Botero-Castro *et al.* 2017), while the *de novo* genome assembly included 94.5% of single-copy orthologs from the Aves data set and 97.7% of the Vertebrata data set as identified by BUSCO (Table 2).

**Table 1:**
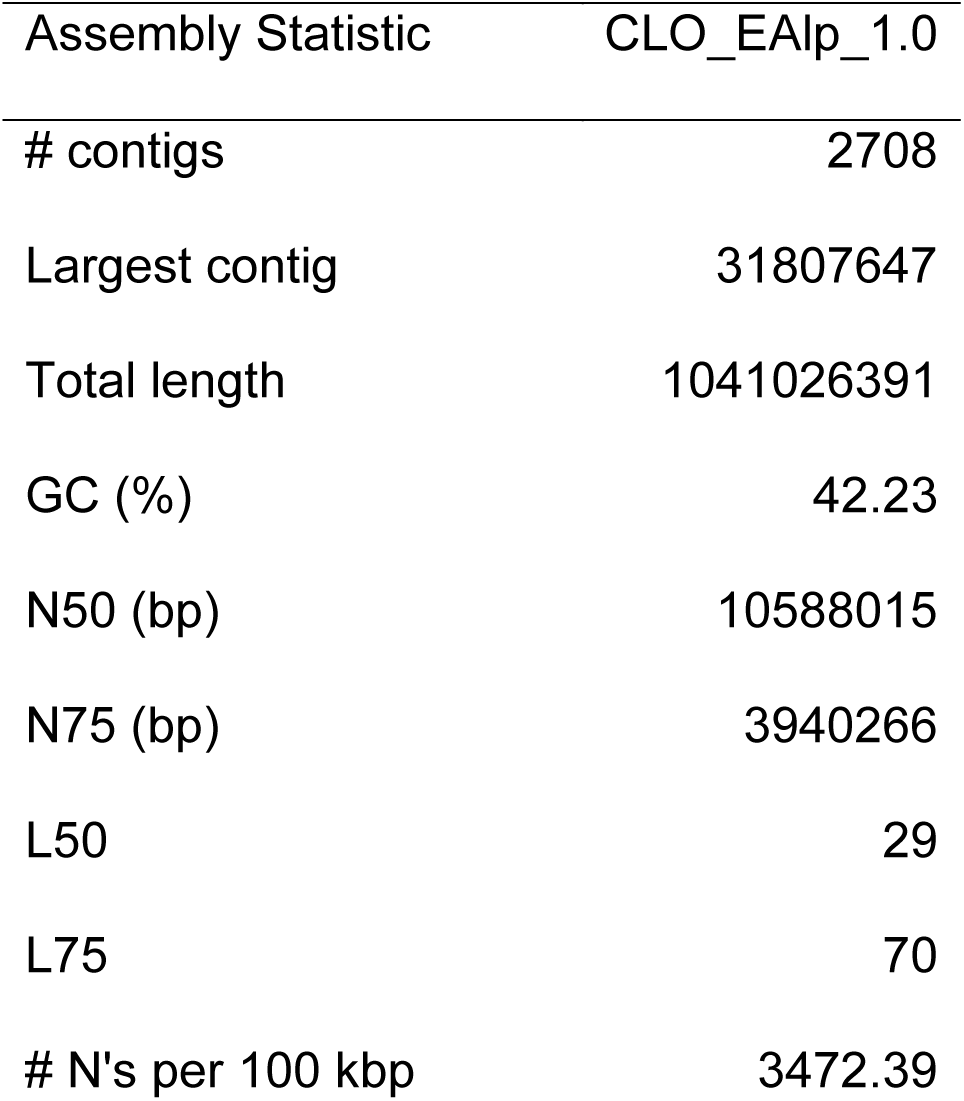
De novo genome assembly metrics estimated using QUAST.

**Table 2:** Output from BUSCO analyses to assess genome completeness by searching for single-copy orthologs from aves and vertebrata datasets.

|  | Aves | Vertebrata |
| --- | --- | --- |
| Complete BUSCOs | 4645 (94.5%) | 2530 (97.7%) |
| Complete and single-copy BUSCOs | 4590 (93.4%) | 2518 (97.4%) |
| Complete and duplicated BUSCOs | 55 (1.1%) | 12 (0.5%) |
| Fragmented BUSCOs | 162 (3.3%) | 36 (1.4%) |
| Missing BUSCOs | 108 (2.2%) | 20 (0.7%) |
| Total BUSCO groups searched | 4915 | 2586 |

We opted not to assemble pseudochromosomes by aligning our *de novo* genome to an existing chromosome-level genome assembly (e.g., Zebra Finch (*Taeniopygia guttata*). While birds generally exhibit strong synteny (Derjusheva *et al.* 2004), avian sex chromosomes and microchromosomes are often comprised of extensive rearrangements (Volker *et al.* 2010). Thus, there is room to improve scaffolds generated in this assembly so that they match full chromosomes through strategies such as Hi-C (Burton *et al.* 2013) or ultra-long read sequencing technology (Ma *et al.* 2018). Functional annotation could also be improved by generating RNA-Seq and protein libraries for larks (Denoeud *et al.* 2008). Thus, while there is room to improve this current assembly, CLO_EAlp_1.0 represents a large step forward toward leveraging the natural history of larks and advanced sequencing technology to further understand avian biology.

## Acknowledgements

We would like to thank Bronwyn Butcher for assistance with lab work. Leonardo Campana gave advice on genome assembly and bioinformatics. Thanks to Diana Carolina Macana for her assistance with field work. This research was supported by a Doctoral Dissertation Improvement Grant from the National Science Foundation (DEB-1601072) to NAM.

## Author Contributions

NAM, PP, and CDC conducted field work. NAM performed lab work, genomic assembly, and analyses. NAM wrote the paper with input from PP, CDC, and IJL. All authors read and approved the final manuscript. The authors declare that they have no competing interests. The funding sponsors had no role in the design of the study; in the collection, analyses, or interpretation of data; in the manuscript writing, and in the publication decision.

